# Astrocyte-derived TNF and glutamate critically modulate microglia reactivity by methamphetamine

**DOI:** 10.1101/2021.02.22.432170

**Authors:** Teresa Canedo, Camila Cabral Portugal, Renato Socodato, Joana Bravo, Tiago Oliveira Almeida, João D. Magalhães, Sónia Guerra-Gomes, João Filipe Oliveira, Nuno Sousa, Ana Magalhães, João Bettencourt Relvas, Teresa Summavielle

## Abstract

Methamphetamine (Meth) is a powerful illicit psychostimulant, widely used for recreational purposes. Besides disrupting the monoaminergic system and promoting oxidative brain damage, Meth also causes neuroinflammation that contributes to synaptic dysfunction and behavioral deficits. Aberrant activation of microglia, the largest myeloid cell population in the brain, is a common feature in neurological disorders linked to cognitive impairment and neuroinflammation. In this study, we investigated the mechanisms underlying the aberrant activation of microglia elicited by Meth in the adult mouse brain. We found that binge Meth exposure caused microgliosis and disrupted risk assessment behavior (a feature that usually occurs in human Meth abusers), both of which required astrocyte-to-microglia crosstalk. Mechanistically, Meth triggered a detrimental increase of glutamate exocytosis from astrocytes (in a manner dependent on TNF production and calcium mobilization), promoting microglial expansion and reactivity. Ablating TNF production or suppressing astrocytic calcium mobilization prevented microglia reactivity and abolished the behavioral phenotype elicited by Meth exposure. Overall, our data indicate that glial crosstalk is critical to relay behavioral alterations caused by acute Meth exposure.

**One Sentence Summary:** Glial crosstalk under methamphetamine exposure

## Introduction

Methamphetamine (Meth) is a potent and highly-addictive psychostimulant that causes long-lasting harmful effects in the central nervous system (CNS)^1, 2^. Meth toxicity is classically characterized by severe disruption of the dopaminergic system, causing oxidative stress and behavioral deficits^3, 4^. More recently, release of proinflammatory mediators and glutamate were also reported^5, 6^.

There is a growing understanding that the interplay between neuronal and glial cells is important for the build-up and maintenance of addiction^7–9^. Gliotransmission is implicated in drug-seeking modulation, with particular focus on glutamatergic signaling^10, 11^, that can trigger calcium influx, leading to reactive oxygen species (ROS) formation and subsequent oxidative damage^12^. However, the overall contribution of such mechanisms to the addictive process remains unclear^13, 14^.

Microglia and astrocytes play crucial roles in brain injury and repair^15, 16^, but their sustained reactivity – often increasing the production of proinflammatory mediators like TNF, glutamate, and ROS^17, 18^ – may result in damage to the brain parenchyma^19, 20^. Under exposure to psychoactive substances, microglia may also become highly reactive, augmenting the release of proinflammatory mediators^13^, and in early abstinence this reactivity might increase the likelihood of relapse^9, 13^. Therefore, a better understanding of the microglia reactivity and associated brain immune-pathways in response to psychostimulants is paramount to implement relevant interventions for treating addictive behaviors. In accordance, we have recently demonstrated that binge alcohol administration to adult mice causes aberrant synaptic pruning and loss of prefrontal cortex excitatory synapses, increasing anxiety-like behavior, which is prevented by pharmacological blockade of Src activation or by reducing TNF production in microglia^21^.

Here, we investigated how Meth interferes with microglia reactivity. Our results showed that the behavioral alterations caused by binge Meth exposure are mediated by astrocyte-microglia crosstalk in which release of glutamate from astrocytes in a TNF/IP3 receptor (IP3R)/SNARE-dependent manner leads to microglial activation, neuroinflammation, and ultimately to changes in behavior.

## Materials and Methods

### Animals

All experiments were in accordance with the Directive 2010/63/EU and approved by the competent authorities Direcção Geral de Alimentação e Veterinária (DGAV) and i3S Animal Ethical Committee (ref.2018-13-TS and DGAV 17469/2012). Researchers involved in animal experimentation were FELASA certified. All efforts were made to minimize animal suffering and the number of animals used.

Mice were housed under specific pathogen-free conditions, controlled environment (20°C, 45–55% humidity) with an inverted 12h/12h light/dark cycle, and allowed free access to food and water. Because of the potential behavioral variability related to the estrous cycle in females^22^, only male mice were used. C57BL/6 male mice were obtained from the i3S animal facility. TNF knockout mice in the C57BL/6 background (referred herein as TNF KO) were kindly supplied by Professor Rui Appelberg (University of Porto). TNF KO mice^21^ were maintained at i3S and genotyped by PCR using ATCCGCGACGTGGAACTGGCAGAA (forward) and CTGCCCGGACTCCGCAAAGTCTAA (reverse) primer pair. IP3R2 KO mice^23, 24^ were held at ICVS animal facility and genotyped PCR using the primer pairs: WT (F, 5’-ACCCTGATGAGGGAAGGTCT-3’; R, 5’-ATCGATTCATAGGGCACACC-3’) and mutant allele (neo-specific primer: F, 5’-AATGGGCTGACCGCTTCCTCGT-3’; R, 5’-TCTGAGAGTGCCTGGCTTTT-3’).

### Mice treatment

Mice were treated using a Meth binge protocol^25, 26^ and randomly assigned to treated group (4×5mg/kg Meth, 2h apart, intraperitoneally) or control (4× isovolumetric saline), and sacrificed 24h after the first administration (**Suppl. Fig. 1A**). Since Meth causes hyperthermia^27^, we controlled body temperature through infrared readings every 20min using subcutaneous tags (Biomark, ID, USA). Meth significantly increased body temperature (**Suppl. Fig. 1B**) but did not exceed critical values. Methamphetamine hydrochloride was imported from Sigma-Aldrich (MO, USA) under special INFARMED license (ref. 290-13).

**Figure 1.**
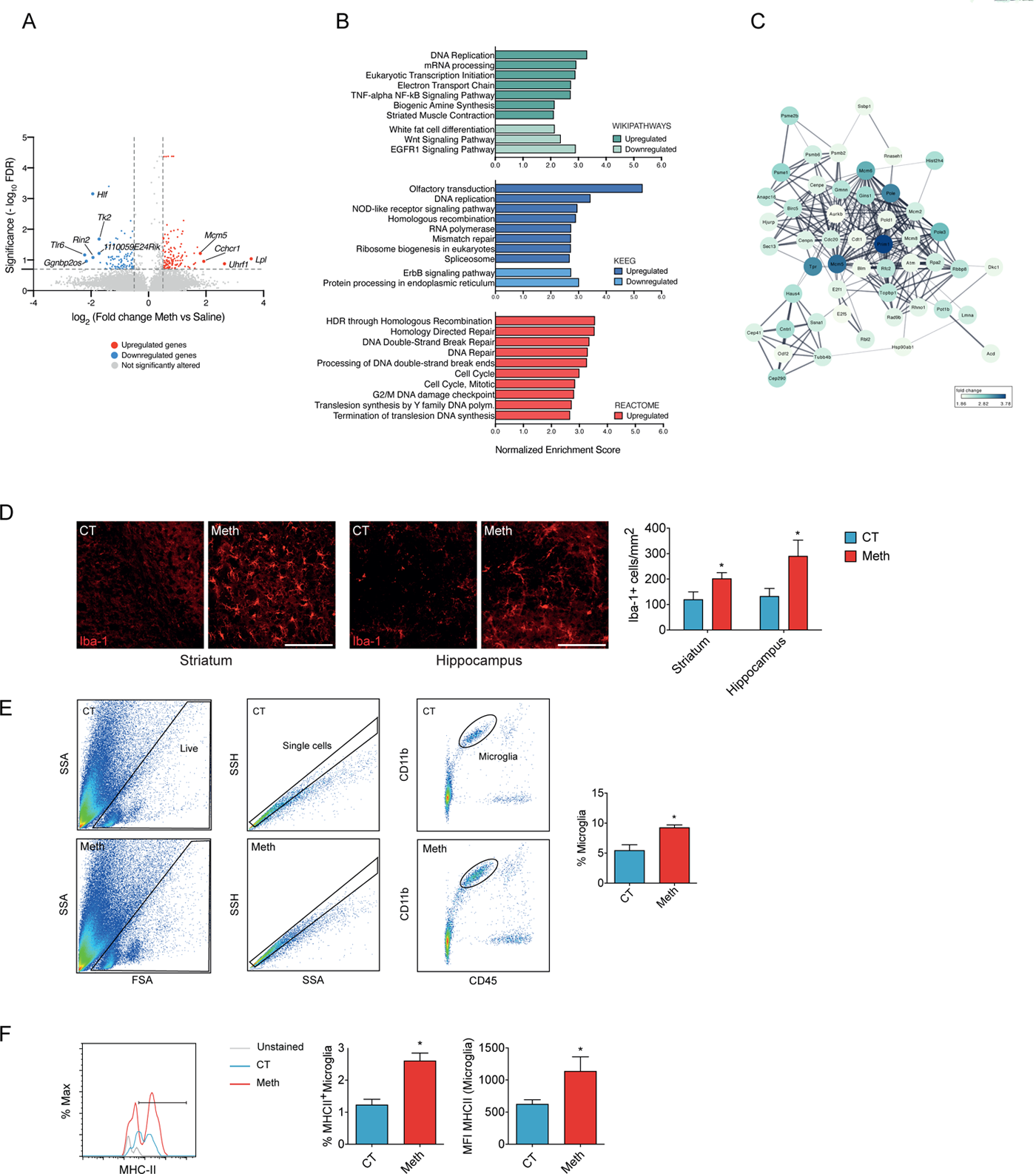
Meth triggers microglial expansion in the brain. **A:** Volcano plot depicting differentially expressed genes of isolated microglia from brains of mice administered with Meth vs Saline (n=3 mice). Non-differentially expressed genes are shown with gray dots, red dots represent significantly upregulated genes and blue dots represent downregulated genes. **B:** Top 10 enriched pathways revealed by Wikiphatways, KEEG and Reactome databases using Gen Set Enrichment Analysis (GSEA). **C:** Network analysis of enriched gene sets involved in cell cycle. Network represents the top of 50 upregulated genes related to cell cycle, upon Meth treatment. **D:** Representative confocal imaging of striatal or hippocampal sections from mice administered with binge Meth or saline (CT) and immunostained for Iba-1. Graphs display the number of Iba1^+^ cells with mean and SEM (3/4 sections *per* animal from n=3 mice). *p<0.05 (unpaired t test). Scale bars, 50μm. **E:** Flow cytometry analyses of microglia cells (CD11b^+^ CD45^Low^) isolated from the brains of mice administered with binge Meth or saline (CT) (n=5 animals for each group). The graph displays the percentage of microglia cells with mean and SEM. *p<0.05 (unpaired t test). **F:** Expression of MHCII by flow cytometry in microglia (CD11b^+^ CD45^Low^) isolated from the brains of mice administered with binge Meth or saline (CT) (n=5 animals for each group). The graph displays the frequency of microglial cells expressing the MHCII marker with mean and SEM. *p<0.05 (unpaired t test).

### Fluorescence-Activated Cell Sorting (FACS) and RNA extraction

Twenty-four hours after methamphetamine administration, animals were perfused under deep anesthesia with ice-cold PBS. The brains were removed and collected in ice-cold medium A (HBBS 1X (Thermo Scientific MA, USA) supplemented with 15mM HEPES and 0.6% glucose both from Sigma-Aldrich (MO, USA). Microglial cells were isolated from adult mice brain exactly as previously described^28^. Microglia (Cd11b^+^, CD45^low^ and CD206^-^) were sorted on the FACS ARIA (BD Immunocytometry Systems, CA, USA) and the RNA was isolated using a RNeasy Plus Micro Kit (Qiagen, Düsseldorf, DE) according to the manufacturer’s instructions. RNA integrity was analyzed using the Bioanalyzer 2100 RNA Pico chips (Agilent Technologies, CA, USA), according to manufacturer instructions.

### Library preparation and Sequencing

Ion Torrent sequencing libraries were prepared according to the AmpliSeq Library prep kit protocol. Briefly, 1ng of highly intact total RNA was reverse transcribed, the resulting cDNA was amplified for 16 cycles by adding PCR Master Mix, and the AmpliSeq mouse transcriptome gene expression primer pool. Amplicons were digested with the proprietary FuPa enzyme, then barcoded adapters were ligated onto the target amplicons. The library amplicons were bound to magnetic beads, and residual reaction components were washed off. Libraries were amplified, re-purified and individually quantified using Agilent TapeStation High Sensitivity tape. Individual libraries were diluted to a 50pM concentration and pooled equally. Emulsion PCR, templating and 550 chip loading was performed with an Ion Chef Instrument (Thermo Scientific MA, USA). Sequencing was performed on an Ion S5XL™ sequencer (Thermo Scientific MA, USA).

### Bioinformatics

Data from the S5 XL run processed using the Ion Torrent platform specific pipeline software Torrent Suite v5.12 to generate sequence reads, trim adapter sequences, filter and remove poor signal reads, and split the reads according to the barcode. FASTQ and/or BAM files were generated using the Torrent Suit plugin FileExporter v5.12. Automated data analysis was done with Torrent Suite™ Software using the Ion AmpliSeq™ RNA plug-in v.5.12 and target region AmpliSeq_Mouse_Transcriptome_V1_Designed.

Raw data was loaded into Transcripotme Analysis Console (4.0 Thremo Fisher Scientific, MA, EUA) and first filtered based on ANOVA eBayes using Limma package, applied to fold changes ≤ −1.5 or ≥1.5 between experimental and control conditions. Significant changes had a p value <0.05 and a false discovery rate <0.2. Genes that significantly downregulated and upregulated by Meth in microglia, following the described criteria, are represented in **Supp. Table 1.**

RNA-seq functional enrichment analysis using Gene Set Enrichment Analysis (GSEA) was performed by WEB-based Gene SeT AnaLysis Toolkit (WebGestalt)^29^. All detectable genes (**Supp. Table 2**) with their corresponding fold-change values were submitted to WebGestalt at http:// http://www.webgestalt.org. GSEA was performed using the open-access available platforms, Wikipathways, KEEG and REACTOME with default settings. Enrichment scores for gene sets were calculated using an FDR cutoff of 0.05 and hypergeometric overlap analysis **(Supp. Table 3)**. Genes retrieved from GSEA datasets were used for constructing a protein-protein interaction network. Such network was generated using Omics Visualizer^30^ and String applications^31^ in Cytoscape.

### Primary cultures

Primary mixed glial cultures were performed as previously described^32, 33^. Briefly, neonatal Wistar rats or C57BL/6 mice were sacrificed, and their cerebral cortices dissected and digested with 0.07% trypsin-EDTA in the presence of DNAse for 15min. Cells were dissociated and seeded in poly-D-lysine-coated T-flasks at 1.5×10^6^ cells/cm^2^ in DMEM GlutaMAX™-I. Culture media was changed every three days up to 21 days. All cultures were kept at 37°C with 95% air/ 5%CO2 in a humidified incubator.

To obtained purified microglia cultures, culture flasks were orbitally shaken (200 rpm, 2h) to detach microglia. Then, culture media containing microglia were collected, centrifuged (453g, 5min), resuspended, and plated in glass coverslips at 2.5×10^5^ cells/cm^2^ in DMEM-F12 GlutaMAX^TM^-I supplemented with 10% FBS, 0.1% Penicillin-Streptomycin and 1ng/ml GM-CSF. Purified microglia were cultured for 4-7 days. Immunolabeling with CD11b showed a purity of 95-99%.

For purified astrocyte cultures, mixed glial cell cultures were shaken to remove non-astrocytic cells. Astrocytes (adherent cells) were detached and split into non-coated T-flasks in DMEM GlutaMAX™-I. Split cultures were re-split at least four times to obtain purified cultures. After that, astrocytes were plated at 2.5×10^4^ cells/cm^2^ in non-coated plates and maintained for 3 to 4 days.

### Astrocyte-conditioned medium and microglia treatment

Astrocytes were seeded at a density of 2.5×10^4^ cells/cm^2^. After two days, cells were left untreated (control) or incubated with 100µM Meth for 24h. Untreated astrocyte-conditioned medium (ACM CT) and conditioned medium from Meth-treated astrocytes (ACM Meth) were collected, centrifuged for debris removal, and frozen at −80°C until used. To evaluate astrocytic conditioned media’s effects, purified microglial cell cultures were exposed to ACM CT or ACM Meth for 24h.

### Flow cytometry

Microglia and macrophages were analyzed, as we previously described^33, 34^. Briefly, mice were anesthetized and perfused with ice-cold PBS. For single-cell suspensions, the whole brain was quickly removed and mechanically homogenized. The cell suspension was passed through a 100μm cell strainer and centrifuged over a discontinuous 70%/30% Percoll gradient. Cells on the interface were collected, pelleted, and resuspended in FACS buffer (2% BSA; 0.1% Sodium Azide in PBS). Cells were counted using the Countess TM automated counter (Thermo Scientific, MA, USA). For microglia and macrophages characterization, the following antibodies were used: CD45-PE (103106), CD11b-Alexa647 (101218), Ly6C-PerCP/Cy5.5 (128012), CCR2-PE/Cy7 (150611), and MHCII-BV421 (107631), all obtained from BioLegend (CA, USA). Samples were evaluated on FACS Canto II (BD Immunocytometry Systems, CA, USA).

### Immunohistochemistry

Mice were anesthetized and perfused with ice-cold PBS, followed by 4% PFA. Brains were post-fixed overnight, cryoprotected using sucrose gradients (15 and 30%), embedded in OCT, frozen and cryosectioned (coronally at 40μm, between Bregma positions 1.0mm-2.0mm) in the CM3050S cryostat (Leica Biosystems, Nussloch, DE). Brain sections were collected on adherent slides and stored at −20°C.

For immunolabeling, brain slides were defrosted and permeabilized with 0.25% Triton X-100 for 15min. Then, brain slices were blocked with 3% BSA, 0.1% Triton X-100 and 5% FBS for 1h. Primary antibodies were incubated overnight (4°C) under the manufacturer’s recommendations. After washing, slices were incubated with corresponding secondary antibodies conjugated to Alexa Fluor for 2h (RT). After PBS washes, sections were mounted using Fluoroshield from Sigma-Aldrich and visualized under a TCS SP5 II confocal microscope (Leica Biosystems). All used antibodies were described in **Suppl. Table 4**.

### Immunocytochemistry

Immunocytochemistry was performed as we previously described^33, 34^. Briefly, after fixation with 4% PFA, cultures were permeabilized with 0.1% Triton X-100 or 10 min and blocked with 3% BSA for 1h. Cells were incubated with primary antibody under the manufacture’s recommendations, washed and incubated with secondary antibodies conjugated with Alexa Fluor 488 or 568 for 1h (RT). Finally, cells were incubated with DAPI, mounted, and visualized using a DMI6000B inverted microscope (Leica Microsystems) with an HCX Plan Apo 63x/1.3 NA glycerol immersion objective. Images were acquired with 4×4 binning using a digital CMOS camera (ORCA-Flash4.0 V2, Hamamatsu Photonics). All antibodies are described in **Suppl. Table 5**.

### Phagocytic assay

Fluorescent latex beads (Sigma-Aldrich) were diluted in a culture medium (0,001%) and incubated for 1h. After that, cells were washed and fixed with 4% PFA. Immunocytochemistry for CD11b was performed, and the phagocytic efficiency of microglia was estimated as described elsewhere with minor modifications^35^.

### Reactive oxygen species determination by fluorescence microscopy

Primary microglia cultures were incubated with CellROX® green reagent from Thermo-Fisher Scientific, according to manufacturer’s recommendations, following PBS washing and fixation with 4% PFA.

### Fluorescent signals quantification and colocalization analysis

For the intensity quantification, images were exported using the Leica LAS AF program in TIFF format (16-bit). Background subtraction of images, image segmentation, and determination of the intensity of the fluorescence signal was processed in FIJI software as before^33^. For colocalization analyses, images were acquired using an HCX Plan Apo 63x/1.4-0.6NA oil immersion objective in 16-bit sequential mode using bidirectional TCS mode at 100Hz with the pinhole kept at one airy in the Leica TCS SP5 II confocal microscope. The Coloc2 plug-in in FIJI was used to establish TNF/GFAP channels’ quantitative colocalization as before^32^.

### Total RNA extraction, cDNA synthesis, and qRT-PCR

From brain tissue, RNA was extracted using the TRIzol^TM^ (Ambion by Life Technologies, MA, USA). RNA from cell cultures was isolated using the RNeasy Mini Kit from Qiagen (Düsseldorf, DE). RNAs quality and concentration were determined using a NanoDrop ND-1000 Spectrophotometer. cDNA synthesis was performed using 1µg of total RNA using RT2 Easy First Strand kit from Qiagen. qRT-PCR was performed using iQ™ SYBR^®^Green Supermix on an iQ™5 multicolor real-time PCR detection system (Bio-Rad, CA, USA). All primers were obtained from Sigma-Aldrich and described in **Suppl. Table 6**. Raw data were analyzed using the ΔΔCT method with Yhwaz serving as the internal control gene and results expressed in relative gene abundance.

### FRET assays

Primary microglia or astrocyte were plated on plastic-bottom culture dishes μ-Dish35mm (iBidi, Martinsried, DE) and transfected with FRET biosensor for glutamate (pDisplay FLIPE-600n, plasmid 13545), ROS (pFRET-HSP33 cys, plasmid 16076) or calcium (pcDNA-D1ER, plasmid 36325), all from Addgene (MA, USA) using jetPRIME^®^ from Polyplus (NY, USA). Imaging was performed using a Leica DMI6000B inverted microscope, and images were processed in FIJI software exactly as before^36^.

### Elevated plus-maze (EPM)

Anxiety-like behavior was assessed using the elevated plus maze (EPM) test precisely as we previously described^21, 37^. The test was conducted in the dark phase of the light/dark cycle. The mice’s movement and location were analysed by an automated tracking system equipped with an infrared-sensitive camera (Smart Video Tracking Software v 2.5, Panlab, Harvard Apparatus). The maze, made of opaque grey polyvinyl, consisted of four arms arranged in a cross-shape; two closed arms have surrounding walls (18cm high), opposing two open arms (all arms 37×6cm). The apparatus was elevated at the height of 50cm. Each mouse was placed on the central platform facing an open arm and allowed to explore the maze for 5min.

### Statistical analysis

A 95% confidence interval was used, and *P* < 0.05 was considered statistically significant. Results were expressed as mean ± SEM (standard error of the mean). Gene clusters were compared by contingency analysis using the Fisher’s exact test and the Baptista-Pike method to calculate the odds-ratio. Experimental units in individual replicates were evaluated for Gaussian distribution using the D’ & Pearson omnibus normality test. When comparing only two experimental groups, the unpaired Student t test with equal variance assumption was used for data with normal distribution, and the Mann-Whitney test was used otherwise. When comparing three or more groups, a one-way analysis of variance (ANOVA), followed by the Bonferroni or Tukey post hoc test was used for data with normal distribution, and the Kruskal-Wallis test followed by Dunn’s multiple comparisons was used otherwise. We used a two-way ANOVA followed by the Sidak test to compare different groups with two independent variables. All quantifications were performed blinded. Statistical analysis was performed using the GraphPad Prism^®^ software version 8.4.3.

## Results

### Microglia exposed to Meth display a core cell cycle-related transcriptomic signature

To clarify the action of Meth in microglia, we used a binge pattern of Meth administration to adult mice (**Suppl. Fig. 1A**) and conducted RNA-Seq analysis in flow cytometry-sorted microglia (CD11b^+^CD45^Low^CD206^-^) from whole brain tissue. Out of 23,930 microglial transcripts identified in the transcriptome dataset, 207 were significantly altered after binge Meth administration (**Fig. 1A** and Suppl. Tables 1 and 2). To pinpoint the most relevant biological pathways altered in the microglial transcriptome after Meth exposure, we performed gene set enriched analysis (GSEA). GSEA using Wikipathways, KEEG, and REACTOME databases revealed a prominent upregulation of cell cycle-related pathways (including DNA Replication, mRNA processing, Eukaryotic Transcription Initiation, Homologous recombination, RNA polymerase, Mismatch repair, DNA Repair, DNA Double-Strand Break, G2/M DNA damage checkpoint, Mitotic Cell Cycle, Cell Cycle) (**Fig. 1B** and detailed data in **Suppl. Table 3**), possibly associated with Meth-induced microglial expansion. Of note, the TNF-alpha NF-kB and the NOD-like receptor signaling pathways, both associated with proinflammatory signaling, were also upregulated (**Fig. 1B**).

The combined cell cycle-related transcriptomic cluster (the top 50 upregulated transcripts are displayed as network in **Fig. 1C**) contained as highest altered transcripts the DNA primase small subunit (Prim1), the DNA polymerase epsilon catalytic subunit A (Pole), the DNA polymerase epsilon subunit 3 (Pole3), the translocated promoter region, nuclear basket protein (Tpr), and the DNA helicases MCM5 and MCM6 (**Fig. 1C**). Thus, initiation of DNA replication, DNA mismatch repair, homologous recombination, and telomere C-strand synthesis (licensed by the epsilon DNA polymerase complex and the MCM complex via 3’-5’ exodeoxyribonuclease and 3’-5’ DNA helicase activities) are plausibly the most strongly microglial pathways affected by Meth exposure.

Next, we compared our cluster of 207 differentially expressed transcripts upon Meth exposure with clusters previously reported for microglial signature program^38, 39^, aging^40^, disease-associated (DAM)^41^, injury-related (IRM)^40^, drug exposure^42, 43^, or the microglial engulfment module^38^ (**Suppl. Fig. 1C and Suppl. Table 7**). Interestingly, we only found a positive association of our Meth-induced cluster with the aging clusters. These data indicate that Meth exposure does not affect the classical signature programs of healthy or diseased microglia but are in line with reports showing that Meth might foster cellular and tissue ageing^44^.

### Meth activates microglia *in vivo*

The increase in expression of cell cycle-related transcripts correlated with a significant increase in the number of Iba-1^+^ cells on tissue sections obtained from the striatum and the hippocampus (**Fig. 1D**) of mice exposed to Meth when compared to saline-treated (control) animals. This increase in microglia numbers was further confirmed using flow cytometry (**Fig. 1E**). We found also an increase in MHC-II expression in microglia (**Fig. 1F**). We also analyzed the brain macrophage population (CD11b^+^CD45^High^) and found no differences between Meth-treated and control mice in total, Ly6C^+^ or Ly6C^+^/CCR2^+^ macrophages (**Suppl. Fig. 2A**). Together, these results indicate that binge Meth administration causes microgliosis.

### Meth activates microglia in an astrocyte dependent-manner

Microglia activation is thought to modify several of their morphological, molecular and functional properties. Therefore, using primary microglia cultures, we investigated whether exposure to Meth altered some of those properties. We found that Meth diminished the microglia capacity to phagocyte inert fluorescent beads (**Fig. 2A**) and did not increase the formation of ROS (**Fig. 2B**) or the expression of iNOS (**Fig. 2C**). We also observed no differences in the mRNA transcript abundance of the proinflammatory cytokines IL-1β, IL-6 and TNF compare to saline-treated microglia (**Fig. 2D**). To further confirm that our microglia cultures were responsive to a classic proinflammatory stimulus, but not to Meth, we treated them with LPS, which as expected increased ROS formation and iNOS expression (**Suppl. Fig. 2B and C**). We also analyzed classic microglial anti-inflammatory markers and found no significant alterations in arginase 1 expression (**Fig. 2E**), nor in the amounts of mRNA transcripts IL-10 and TGFβ (**Fig. 2F**). We concluded that Meth does not activate microglia in a cell-autonomous manner and that the transcriptomic changes associated with microgliosis observed in vivo might result from crosstalk between microglia and other cell types.

**Figure 2.**
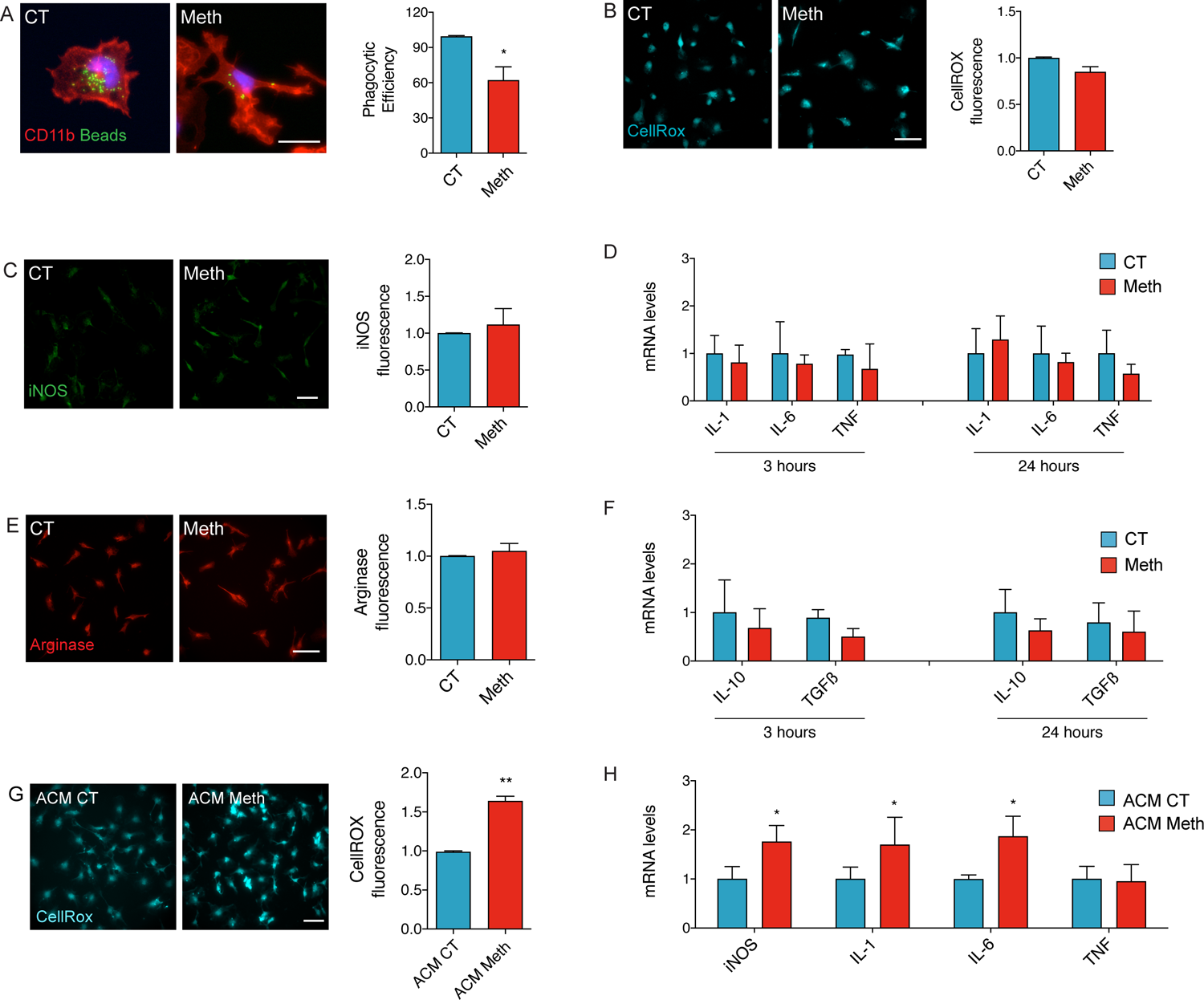
Microglia activation triggered by Meth requires Astrocytes. **A:** Fluorescence imaging of CD11b (red) in primary cortical microglia incubated with microbeads (green) and treated with 100μM Meth for 24h (n=3 independent cultures). Graph (means and SEM) displays phagocytic efficiency. *p<0.05 (unpaired t test). Scale bar, 10μm. **B:** Fluorescence imaging of primary cortical microglia incubated with the CellRox^®^ green reagent and treated with 100μM Meth for 24h (n=3 independent cultures). Graph (means and SEM) displays the CellRox^®^ intensity normalized to the Control values (unpaired t test). Scale bar, 10μm. **C:** Fluorescence imaging of primary cortical microglia immunolabeled for iNOS (green) treated with 100μM Meth for 24h (n=3 independent cultures). Graph (means and SEM) displays iNOS intensity normalized to the Control values (unpaired t test). Scale bar, 10μm. **D:** qRT-PCR for IL-1β, IL-6 or TNF from primary cortical microglia treated with 100μM Meth for 3h or 24h (n=3 independent cultures). Graphs (means and SEM) display the indicated transcripts’ mRNA expression levels (unpaired t test). **E:** Fluorescence imaging of arginase in primary cortical microglia treated with 100μM Meth for 24h (n=3 independent cultures). Graph (means and SEM) displays arginase intensity normalized to the CT values (unpaired t test). Scale bar, 10μm. **F:** qRT-PCR for IL-10 or TGFβ from primary cortical microglia treated with 100μM Meth for 3h or 24h (n=3 independent cultures). Graphs (means and SEM) display the indicated transcripts’ mRNA expression levels (unpaired t test). **G:** Fluorescence imaging of primary cortical microglia (n=3 independent cultures) incubated with the CellRox^®^ green reagent and then exposed to conditioned media from primary cortical astrocytes (ACM) treated with 100μM Meth or not (CT). Graph (means and SEM) displays the CellRox intensity normalized to the ACM CT values. *p<0.05 (unpaired t test). Scale bar, 10μm. **H:** qRT-PCR for iNOS (C), IL-1β (D), IL-6 (E), or TNF (F) from primary cortical microglia exposed to ACM CT or ACM Meth for 24h (n=3-5 independent cultures). Graphs (means and SEM) display the mRNA fold change for the indicated transcripts.

Because astrocyte-derived signaling is essential in microglia activation^45^, we tested the hypothesis that astrocytes could mediate Meth-induced microglia activation. To do that, we exposed primary cortical microglia to conditioned media (CM) obtained from primary cortical astrocytes treated with Meth (ACM Meth) or CM from control astrocyte cultures (ACM CT). Neither Meth nor ACM Meth affected astrocytic or microglial viability (**Suppl. Fig. 2D-F**). Using the CellRox green reagent, we found an increase in ROS production in primary cortical microglia exposed to ACM Meth compared with cultures exposed to ACM CT (**Fig. 2G**). Using the FRET HSP biosensor^46^, we observed a consistent and fast increase (within 5 min) of ROS generation in living primary microglia exposed to ACM Meth (**Suppl. Fig. 3A**). Besides, primary cortical microglia treated with ACM Meth displayed higher mRNA levels of the proinflammatory markers iNOS, IL-1β, and IL-6, but not TNF (**Fig. 2H**). Primary cortical microglia exposed to ACM Meth also displayed enhanced iNOS expression compared with cultures incubated with ACM CT (**Suppl. Fig. 3B**). We concluded that upon Meth exposure, astrocytes could induce microglial activation.

<u>Meth causes glutamate release via TNF and IP_3_-dependent</u> Ca^2+^ mobilization in astrocytes Astrocytes are critical players in regulating neuroinflammation^47^. Of note, our RNA-Seq data revealed a Meth-induced enrichment of gene transcripts associated with the TNF-alpha NF-kB Signaling Pathway (**Fig. 1B**). Besides, TNF has emerged as an essential mediator of brain homeostasis^48^. We observed increased TNF expression in specific brain regions following Meth exposure (**Suppl. Fig 3D**), which was also previously reported^49^. The secretion of high amounts of TNF activates TNF receptor 1 and leads to a massive release of glutamate from astrocytes^50^. Accordingly, we observed by double-labeling immunofluorescence an increase in TNF content in astrocytes (GFAP^+^ cells) in the hippocampus of mice exposed to Meth (**Fig. 3A**), and using the glutamate-release FRET biosensor FLIPE600n^SURFACE 51^, we found that TNF promoted a fast and sustained release of glutamate from living cortical astrocytes (**Suppl. Fig. 3C**). In addition, Meth also caused robust glutamate release in cortical astrocytes from WT mice (**Fig. 3B**). However, Meth was inefficient in triggering glutamate release in cortical astrocytes from TNF-deficient mice (**Fig. 3B**), confirming that autocrine TNF signaling plays a crucial role in Meth-induced glutamate release from astrocytes.

**Figure 3.**
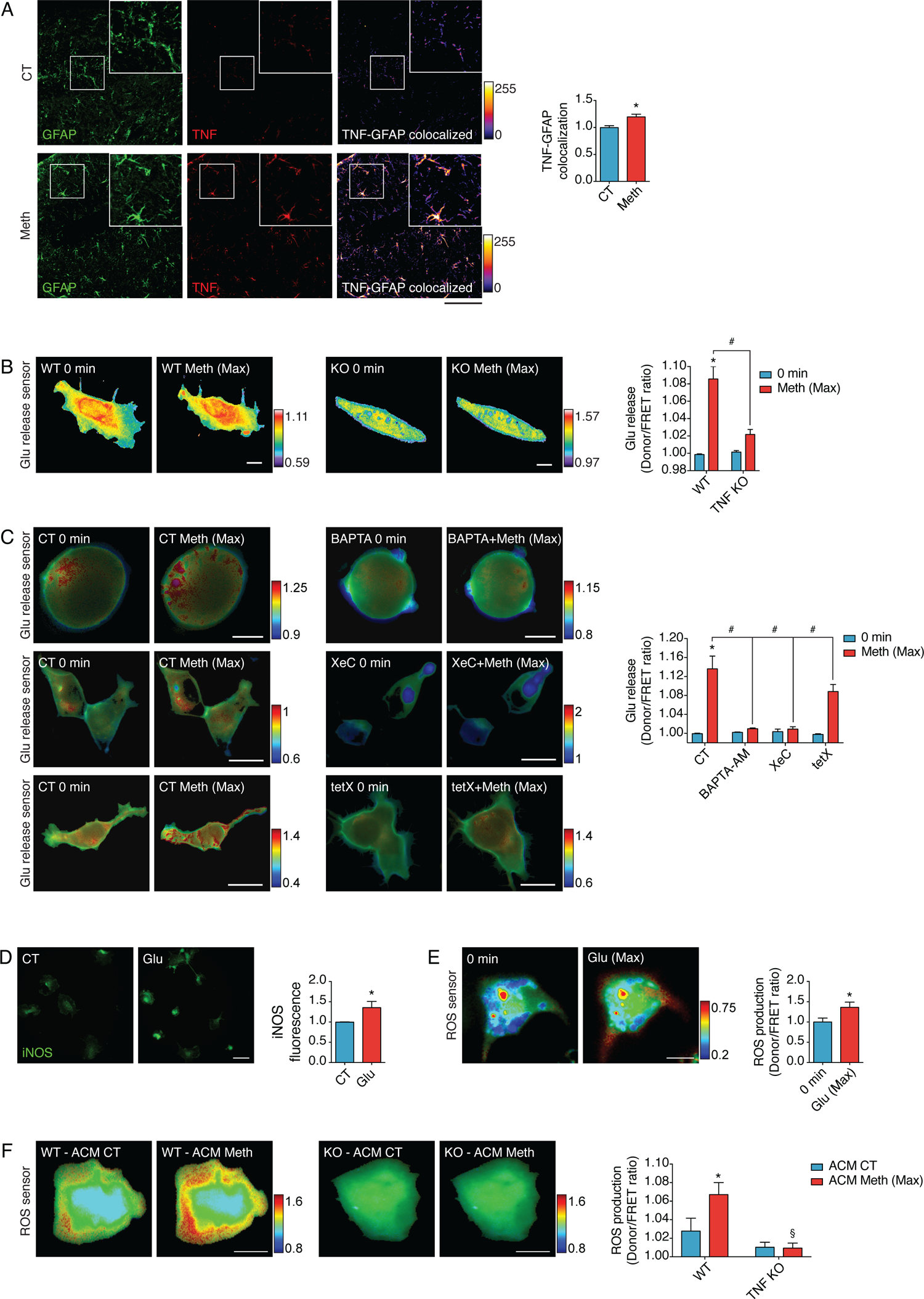
Meth activates microglia via astrocytic TNF production. **A:** Confocal imaging of hippocampal sections from mice treated with Meth or saline (CT) and immunostained for GFAP (green) and TNF (red). Graphs display the GFAP/TNF colocalization puncta (upper graph) or GFAP intensity (bottom graph) normalized to the CT values (3/4 sections *per* animal from n=3 mice). *p<0.05 (unpaired t test). Scale bars, 50μm **B:** Primary cortical astrocytes from WT or TNF KO mice expressing the glutamate release FRET biosensor (FLIPE) were exposed to Meth 100µM. Time-lapses of CFP/FRET ratio changes for the FLIPE biosensor (normalized at 0 min) shows the maximum effect of Meth in both genotypes and are coded according to the scale (n=3-8 cells pooled across 2-3 independent experiments). Scale bars, 10μm **C:** Primary cortical astrocytes expressing the glutamate release FRET biosensor (FLIPE) were exposed to Meth, BAPTA-AM (10µM) + Meth 100µM (upper panels), XestosponginC (XeC; 500nM) + Meth 100µM (middle panels) or Tetanus toxin (Tet; 500nM) + Meth (bottom panels). Time-lapses of CFP/FRET ratio changes for the FLIPE biosensor (normalized at 0 min) show the maximum effect of Meth and are coded according to the scale (n=5-7 cells pooled across 3 independent experiments). *p<0.05 (two-way ANOVA vs CT 0 min); # p<0.05 (two-way ANOVA vs CT Meth). Scale bars, 10μm. **D:** Fluorescence imaging of primary cortical microglia immunolabeled for iNOS and treated with glutamate 100µM (n=3 independent cultures). Graph (means and SEM) displays iNOS intensity normalized to the CT. *p<0.05 (Mann-Whitney test). Scale bar, 10μm. **E:** Primary cortical microglia expressing the ROS FRET biosensor (HSP) were exposed to glutamate 100µM. Time-lapses of CFP/FRET ratio changes for the HSP biosensor (normalized at 0 min) shows the maximum effect of Meth and are coded according to the scale (n=5 cells pooled across two independent experiments). *p<0.05. Scale bars, 10μm. **F:** Primary cortical microglia from WT or TNF KO mice expressing the ROS FRET biosensor HSP were incubated with ACM CT and then exposed to ACM Meth 100µM. Time-lapses of CFP/FRET ratio changes for the HSP biosensor (normalized at 0 min) show the maximum effect of Meth and are coded according to the scale (n=4 cells pooled across two independent experiments). *p<0.05, §non-significant. Scale bars, 10μm.

Astrocytes can release glutamate from intracellular pools through various mechanisms, including Ca^2+^-dependent and -independent pathways^52^. To test whether glutamate release from astrocytes under Meth exposure is Ca^2+^-dependent, we chelated cytosolic Ca^2+^ with BAPTA-AM and observed an inhibition of Meth-induced glutamate release (**Fig. 3C**), suggesting that elevation of cytosolic Ca^2+^ is necessary for Meth-triggered astrocytic glutamate release.

The rise in cytosolic Ca^2+^ required for glutamate release from astrocytes may originate from the endoplasmic reticulum (ER) through the Ca^2+^-release channel inositol triphosphate receptor (IP3R)^53^. Using the D1ER FRET biosensor^54^, which detects the efflux of Ca^2+^ from the ER into the cytosol, we monitored the mobilization of Ca^2+^ in living astrocytes exposed to Meth or TNF (**Suppl. Fig. 3E**). Treatment of primary cortical astrocytes with Meth (**Suppl. Fig. 3E, blue circles**) or TNF (**Suppl. Fig. 3E, red circles**) triggered a fast and sustained decrease in the FRET/CFP ratio of the D1ER biosensor, indicating that both Meth and TNF promoted the mobilization of Ca^2+^ from the ER to the cytosol. To investigate the role of IP3R in Meth-induced Ca^2+^-mobilization, we used Xestospongin C (XeC)^55^, an IP3R antagonist. We observed that XeC abolished glutamate release in living primary astrocyte cultures exposed to Meth (**Fig. 3C**) or TNF (**Suppl. Fig. 3F**), and concluded that IP3R-dependent Ca^2+^ mobilization is involved in Meth-induced glutamate release.

To test whether in Meth-treated astrocytes, glutamate was released through an exocytic mechanism^56^, we used the tetanus toxin to prevent Ca^2+^-dependent assembling of the dnSNARE complex and the fusion of exocytic vesicles with the membrane^57^. In these conditions, we observed a large attenuation in the Meth-induced CFP/FRET ratio change of the FLIPE biosensor (**Fig. 3C**), indicating that, in astrocytes, Meth stimulates the exocytosis of glutamate-containing vesicles in a Ca^2+^-dependent manner.

Because TNF controls astrocytic glutamate release, we hypothesized that TNF/glutamate signaling might be directly involved in microglia activation by astrocytes that were exposed to Meth. Accordingly, we found that treating primary microglia with glutamate increased iNOS expression (**Fig. 3D**). Glutamate treatment also promoted fast and sustained ROS generation in living primary cortical microglia as revealed by using the FRET HSP ROS biosensor (**Fig. 3E**). While the CM obtained from WT astrocytes exposed to Meth promoted ROS generation in primary microglia (**Fig. 3F**), the CM obtained from TNF-deficient astrocytes exposed to Meth failed to increase microglial ROS production (**Fig. 3F**), confirming that TNF/glutamate signaling is necessary to induce microglial activation by astrocytes.

<u>TNF and</u> IP3R2<u>-dependent</u> Ca^2+^ mobilization are required for microglia activation in vivo Because Meth activates microglia *via* TNF-to-IP3R signaling in astrocytes, we evaluated whether Meth-induced microgliosis required this signaling *in vivo*. Knowing that the IP3R isoform 2 is the primary IP3 receptor in astrocytes and the major source of Ca^2+^-translocation from the ER into the cytosol in these cells^58^, we challenged IP3R2 KO, and TNF KO mice with binge Meth administration (as described in Suppl. Fig. 1A). We observed that the Meth-induced microgliosis in the striatum and in the hippocampus was prevented in both KO mice compared to WT (**Fig. 4A**). Consistently with these findings, flow cytometry showed that the Meth-induced increase in the microglia population was also prevented in TNF KO and IP3R2 KO mice (**Fig. 4B**).

**Figure 4.**
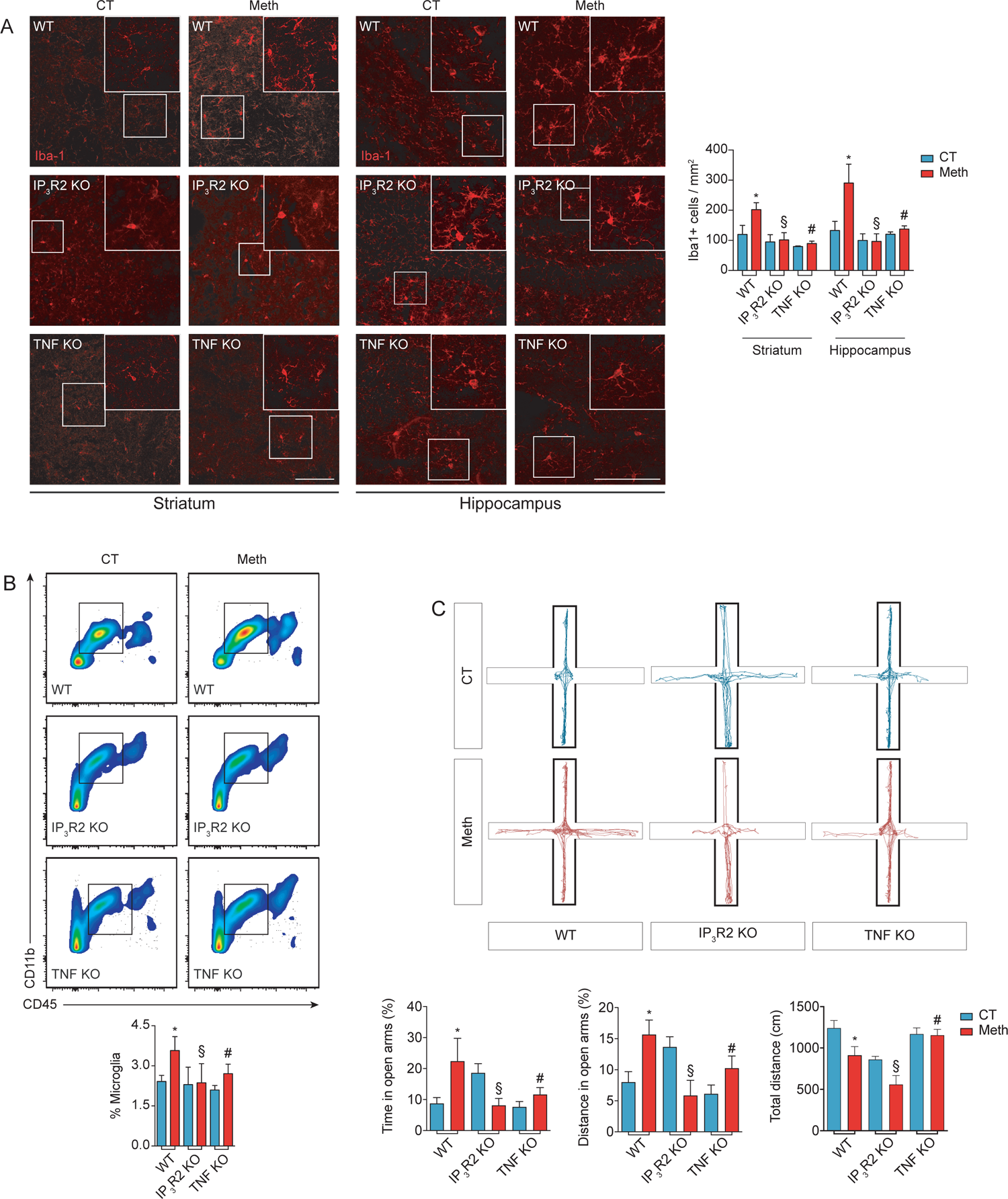
TNF or IP_3_R2 deficiency prevents Meth-induced microgliosis and behavioral changes. **A:** Confocal imaging of striatal or hippocampal sections from WT, IP_3_R2 KO, or TNF KO mice administered with binge Meth (3/4 sections *per* animal from n=3 mice) or saline (CT; n=3) and immunostained for Iba-1. Graphs (means and SEM) display the number of Iba-1+ cells *p<0.001 WT-CT vs. WT-Meth; §non-significant (IP_3_R2 KO-CT vs. IP_3_R2 KO-Meth and ^#^non-significant TNF KO-CT vs. TNF KO-Meth). Two-way ANOVA with the Sidak post hoc analysis. Scale bars, 50μm. **B:** Flow cytometry analysis of microglia cells (CD11b+ CD45Low) isolated from WT, IP3R2 KO, or TNF KO mice injected with Meth or saline (CT) (n=5-9 animals per group). The graph displays the percentage of microglia cells with mean and SEM. *p<0.05 WT-CT vs. WT-Meth; ^§^non-significant IP_3_R2 KO-CT vs. IP_3_R2 KO-Meth and ^#^non-significant TNF KO-CT vs. TNF KO-Meth. Two-way ANOVA with Fisher’s LSD post hoc analysis. **C:** WT, IP_3_R2 KO, and TNF KO animals were evaluated in the EPM 24 hours after being a binge pattern of Meth or saline (CT) administration (n=6-13 animals). CT and Meth-treated mice displayed significant differences in the time spent in the open arms (OA) in the distance traveled in the OA, and in total distance traveled. Graphs display means and SEM. *p<0.05, WT-CT vs. WT-Meth; ^§^non-significant IP_3_R2 KO-CT vs. IP_3_R2 KO-Meth and ^#^non-significant TNF KO-CT vs. TNF KO-Meth. Two-way ANOVA with the Sidak post hoc analysis.

Excessive glutamate and microglia overactivation can negatively affect behavior^59^. Because Meth-induced TNF production led to glutamate release from astrocytes in an IP3R-dependent manner and activated microglia, we hypothesized that blocking TNF or IP3R signaling could prevent the behavioral alteration elicited by Meth. When tested in EPM, WT mice exposed to Meth displayed increased time and distance traveled in the open arms (**Fig. 4C**) and decreased frequency of stretch-attended postures (**Suppl. Fig. 4**), while the total traveled distance was lower than for the saline group (**Fig. 4C**). This behavioral pattern, which is consistent with decreased risk assessment and typical of psychostimulant intake, and was significantly attenuated in TNF or IP3R2 KO mice (**Fig. 4C**). These *in vivo* data confirm the relevance of the the TNF/Ca^2+^ mobilization-signaling for Meth-induced microgliosis and behavioral effects.

## Discussion

Although it was previously observed that Meth induces a microglia proinflammatory response *in vivo*^25, 60^, the mechanisms involved in this process are still poorly understood. We found that Meth-induced microglia reactivity requires a crosstalk with astrocytes, mediated by glutamate release in a TNF- and IP3R/Ca^2+^-dependent manner and that blocking TNF-signaling prevented both microgliosis and the loss of risk assessment behavior elicited by Meth.

Consistently with previous findings^61, 62^, our study shows that binge Meth caused microglial expansion and increased the expression of proinflammatory markers that are hallmarks of many neurodegenerative diseases^63^. The range of enriched pathways related to cell cycle modulation that associate with microglial expansion, confirms the relevance of this Meth-induced effect. To characterize the molecular mechanisms involved in Meth-induced microglia activation, we analysed Meth effects directly on purified microglia cultures. In contrast with a previous work reporting that Meth induces a proinflammatory response in an immortalized microglial cell line^64^, our results demonstrated that Meth does not directly induce a proinflammatory phenotype in primary microglia. Nonetheless, and corroborating our findings, Frank and colleagues observed that Meth fails to induce the expression of proinflammatory cytokines in microglial cultures despite up-regulating IL-1, IL-6, and TNF *in vivo*^65^. Likewise, our primary microglia cultures were highly responsive to LPS, excluding the possibility that the lack of a direct Meth effect could be due to microglia anergy^65^. Similarly, cocaine was reported to be ineffective in directly inducing the expression of microglial TNF mRNA levels^66^ *in vitro*.

Because Meth activated microglia *in vivo*, we tested the hypothesis that this activation could result from an interplay with other cell types. Reactive astrocytes^67^ are observed in several models of Meth exposure^68–70^, including human cerebral organoids^71^, and persistently associated with increased neurotoxicity and neuroinflammation, strengthening the likelihood of an astrocyte-mediated microglial response. Astrocytes seem to control immune activation *via* secretion of multiple molecular factors^72, 73^. Among them, TNF emerged as an essential mediator of brain homeostasis^48^. Increased. We demonstrated that Meth increased TNF content in hippocampal astrocytes *in vivo* and *in vitro*, suggesting that TNF may play an important role in microglia activation by Meth-sensitized astrocytes. Indeed, it has been reported that an autocrine/paracrine TNF-dependent TNF receptor 1 activation promotes glutamate release from astrocytes^50^, while TNF inhibitors strongly reduce glutamate release in cultured astrocytes^74^. In line with this, we also observed that while Meth triggered rapid and sustained glutamate release from astrocytes obtained from wild-type mice, it failed to do so in astrocytes obtained from TNF-deficient mice. In addition, TNF downregulates the glutamate transporter EAAT-2 on astrocytes, compromising glutamate clearance from the extracellular space, which contributes to an hyperglutamate state and promotes excitotoxic glutamate signaling^75, 76^. Excitotoxicity associates positively with the progression of several neurodegenerative diseases^77^. Meth, by acting on the trace amine-associated receptor 1 (TAAR1), induces excitotoxicity through downregulation of EAAT-2 transcription and activity in astrocytes^78^. In this context, our results strongly suggest that glutamate is a critical modulator in Meth-induced microglial activation. Corroborating this hypothesis, we observed that Meth failed to induce microgliosis and loss of risk-assessment behavior in TNF-deficient mice. Interestingly, TNF-deficient mice were previously reported to self-administer more Meth^79^, which according to our data, may also result from reduced astrocyte-microglia reactivity, and not only from increased dopamine availability, as previously suggested^26^.

Astrocytes release glutamate through different pathways, including Ca^2+^-dependent and -independent mechanisms^52^. The ER serves as a major source for astrocytic mobilization of intracellular Ca^2+^ *via* IP3R^12, 80^. We evaluated the involvement of IP3 in Meth-induced glutamate release from astrocytes and confirmed that it occurs in an IP3-dependent way. Accordingly, when we administered Meth to IP3R2-deficient mice, microgliosis and behavioral changes were prevented, suggesting that astrocytic IP3R/Ca^2+^ signaling is required for microglia activation triggered by Meth.

Astrocytes were also recently demonstrated as critical modulators of the reward system, responding to amphetamine-elicited dopaminergic signaling and regulating excitatory neurotransmission through ATP/adenosine activation of neuronal A1 adenosine receptors^81^. Our results provide further mechanistic insight reinforcing the astrocytes’ role in reward and addiction by regulating microglial reactivity.

Collectively, our findings show that astrocytes cause the activation of microglia in acute Meth-exposure *via* glutamate release in a TNF/IP3R2-Ca^2+^-dependent manner (**Fig. 5**), leading to behavioural alterations. Comprehending how microglial reactivity and neuroinflammation will adapt throughout prolonged exposure to Meth, particularly during withdrawal, will further increase the translational significance of our findings and contribute to identifying novel molecular targets with therapeutic value in psychostimulant abuse.

**Figure 5.**
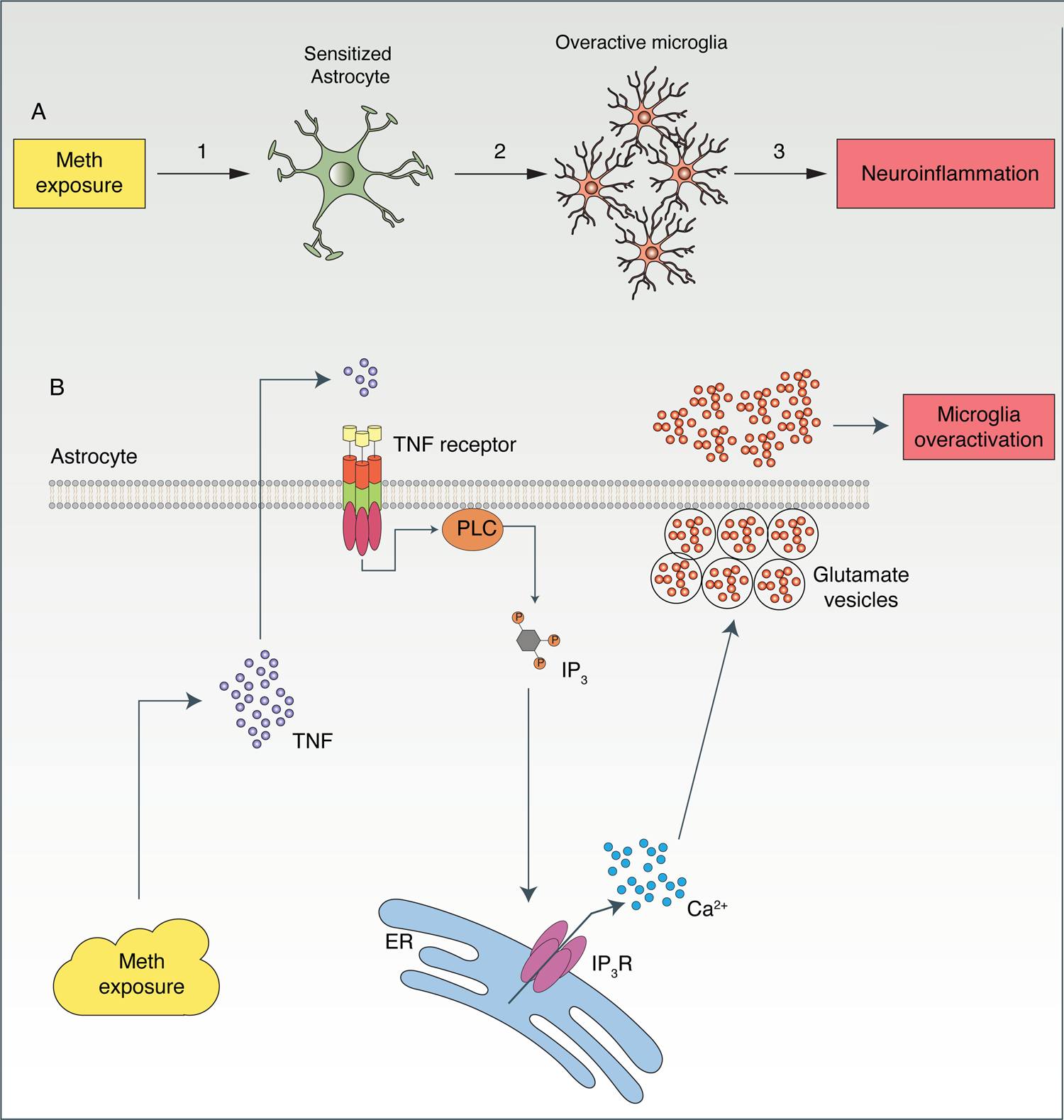
Meth-induced microglia activation occurs via astrocytes. **A:** Exposure to Meth induces astrocytic sensitization (1). Meth-sensitized astrocytes secrete soluble factors (2) that will act on microglia cells, inducing their activation. **B:** In astrocytes, Meth triggers the production (1) and secretion (2) of TNF. TNF acts on astrocytic TNF receptors in an autocrine/paracrine manner, leading to the activation of PLC (3). TNF-induced PLC activation produces the second messenger IP_3_(4) that interacts with IP_3_ receptors on the ER (5). Activation of IP_3_R2 promotes Ca^2+^-mobilization from the ER into the cytosol (6), consequently increasing glutamate release (7). Increased glutamate and TNF content in the extracellular milieu promotes the activation of microglia (8). TNF: Tumor necrosis factor; PLC: Phospholipase C; IP_3_: Inositol (1,3,4) phosphate; ER: Endoplasmic reticulum; Ca^2+^: Calcium ions.

### Funding and Disclosure

This work was financed by FEDER - Fundo Europeu de Desenvolvimento Regional funds through the COMPETE 2020 - Operational Programme for Competitiveness and Internationalisation (POCI), Portugal 2020, and by Portuguese funds through FCT - Fundação para a Ciência e a Tecnologia/Ministério da Ciência (FCT), Tecnologia e Ensino Superior in the framework of the project POCI-01-0145-FEDER-030647 (PTDC/SAU-TOX/30647/2017) in TS lab. FEDER Portugal (Norte-01-0145-FEDER-000008000008—Porto Neurosciences and Neurologic Disease Research Initiative at I3S, supported by Norte Portugal Regional Operational Programme (NORTE 2020), under the PORTUGAL 2020 Partnership Agreement, through the European Regional Development Fund (ERDF); FCOMP-01-0124-FEDER-021333). CCP and RS hold employment contracts financed by national funds through FCT – in the context of the program-contract described in paragraphs 4, 5, and 6 of art. 23 of Law no. 57/2016, of August 29, as amended by Law no. 57/2017 of July 2019. TC and AM were supported by FCT (SFRH/BD/117148/2016 and IF/00753/2014). Work in JBR lab was supported by the FCT project PTDC/ MED-NEU/31318/2017. JFO was also supported by FCT projects PTDC/MED-NEU/31417/2017 and POCI-01-0145-FEDER-016818; Bial Foundation Grants 207/14 and 037/18, by National funds, through FCT - project UIDB/50026/2020 and UIDP/50026/2020; and by the projects NORTE-01-0145-FEDER-000013 and NORTE-01-0145-FEDER-000023, supported by Norte Portugal Regional Operational Programme (NORTE 2020), under the PORTUGAL 2020 Partnership Agreement, through the European Regional Development Fund (ERDF).

We acknowledge the support of the following i3S Scientific Platforms: Animal Facility, Cell Culture and Genotyping (CCGen), Translational Cytometry Unit (TraCy), and Advanced Light Microscopy (ALM), a member of the national infrastructure PPBI-Portuguese Platform of BioImaging (POCI-01–0145-FEDER-022122). The RNAseq technique was performed at the Genomics i3S Scientific Platform with the assistance of Mafalda Rocha as a result of the GenomePT project (POCI-01-0145-FEDER-022184), supported by COMPETE 2020 - Operational Programme for Competitiveness and Internationalization (POCI), Lisboa Portugal Regional Operational Programme (Lisboa2020), Algarve Portugal Regional Operational Programme (CRESC Algarve2020), under the PORTUGAL 2020 Partnership Agreement, through the European Regional Development Fund (ERDF), and by FCT. We also acknowledge Rui Applelberg for generously supplying TNF KO mice, and Maria Summavielle for her contribution in assembling the figures illustrating this publication.

---

**Supplementary Figure 1.**
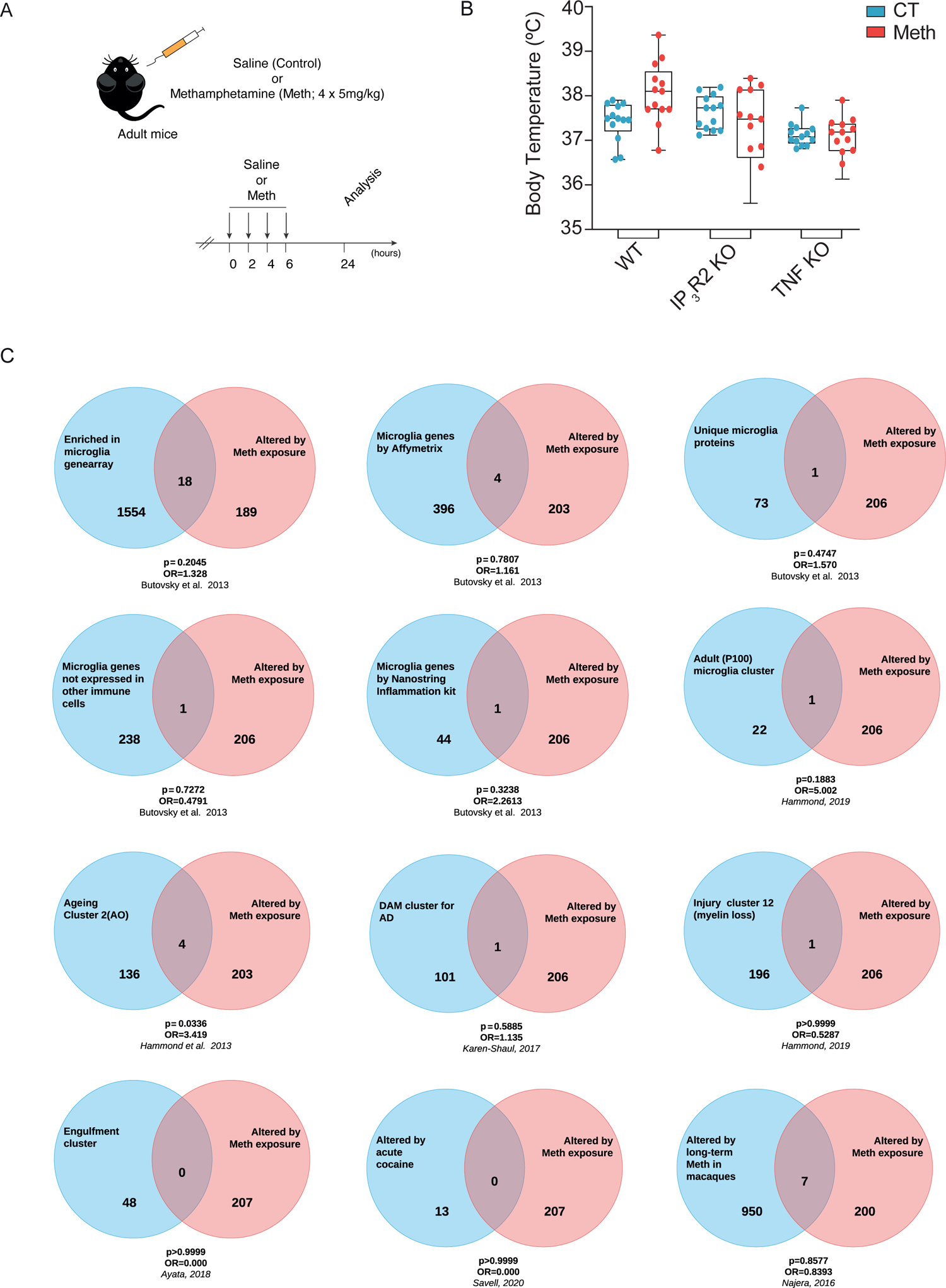
**A**: Schematic representation of binge Meth administration. **B:** WT, IP_3_R2 KO, and TNF KO mice were administered saline (CT) or Meth. The whisker plots represent the median (line within the box), maximum (top whisker) and minimum (bottom whisker) values of mice’s body temperature during the Meth administration protocol. Temperatures were evaluated at 13 time points, each point represents the mean temperature (n=3 animals *per* group) for one timepoint. **C:** Venn’s diagrams representing cluster analysis comparing the 207 Meth-altered genes cluster found in our RNA.-seq analysis, with clusters previously reported for healthy^38, 39^, aging^40^, disease-associated (DAM)^41^, injured^40^, drug exposed microglia^42, 43^, or with clusters previously associated to specific microglia functions^38^. Comparisons were conducted by contingency analysis, using the Fisher’s exact test and the Baptista-Pike method to calculate the odds-ratio. Significance was set at p<0.05. A comprehensive list of the shared genes in each case is available in **Suppl. Table 7**.

**Supplementary Figure 2.**
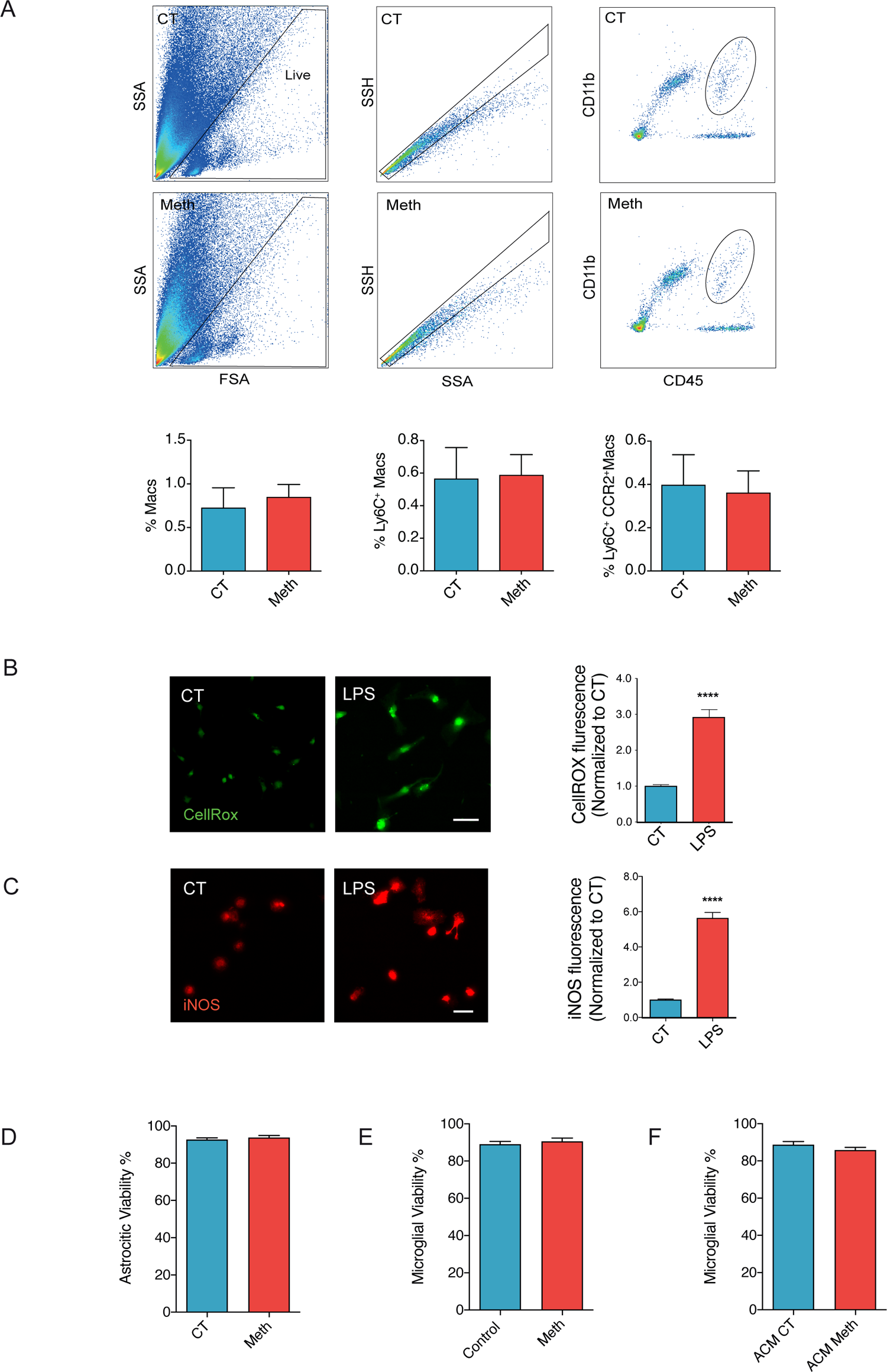
**A:** Flow cytometry analyses macrophages (CD11b+ CD45high) isolated from the brains of mice injected with Meth or saline (CT) (n=5 animals for each group). Graphs display with mean and SEM of the percentage of macrophages, the percentage of macrophages expressing activation markers such as Ly6C+ and Ly6C+CCR2+. **B:** Primary cortical microglia cells incubated with the CellRox green reagent and treated with 1µM LPS (n=3 different cultures). Graph (means and SEM) displays the CellRox intensity normalized to the control values (unpaired t test). Scale bar, 10μm. **C:** Fluorescence imaging of primary cortical microglia immunolabeled for iNOS treated with 1µM LPS (n=3 independent cultures). Graph (means and SEM) displays iNOS intensity normalized to the Control values (unpaired t test). Scale bar, 10μm. **D:** Viability of astrocytes were examined by Hoechst staining under 100μM Meth. Graph represent (means and SEM) the percentage of cell viability upon Meth exposure compared to control (CT) condition. **E:** Viability of microglial cells were examined by Hoechst staining under 100μM Meth. Graph represent (means and SEM) the percentage of cell viability upon Meth exposure compared to control (CT) condition. **F:** Viability of microglia were examined by Hoechst staining under ACM Meth exposure. Graph represent (means and SEM) the percentage cell viability upon ACM Meth exposure compared to control condition (ACM CT).

**Supplementary Figure 3.**
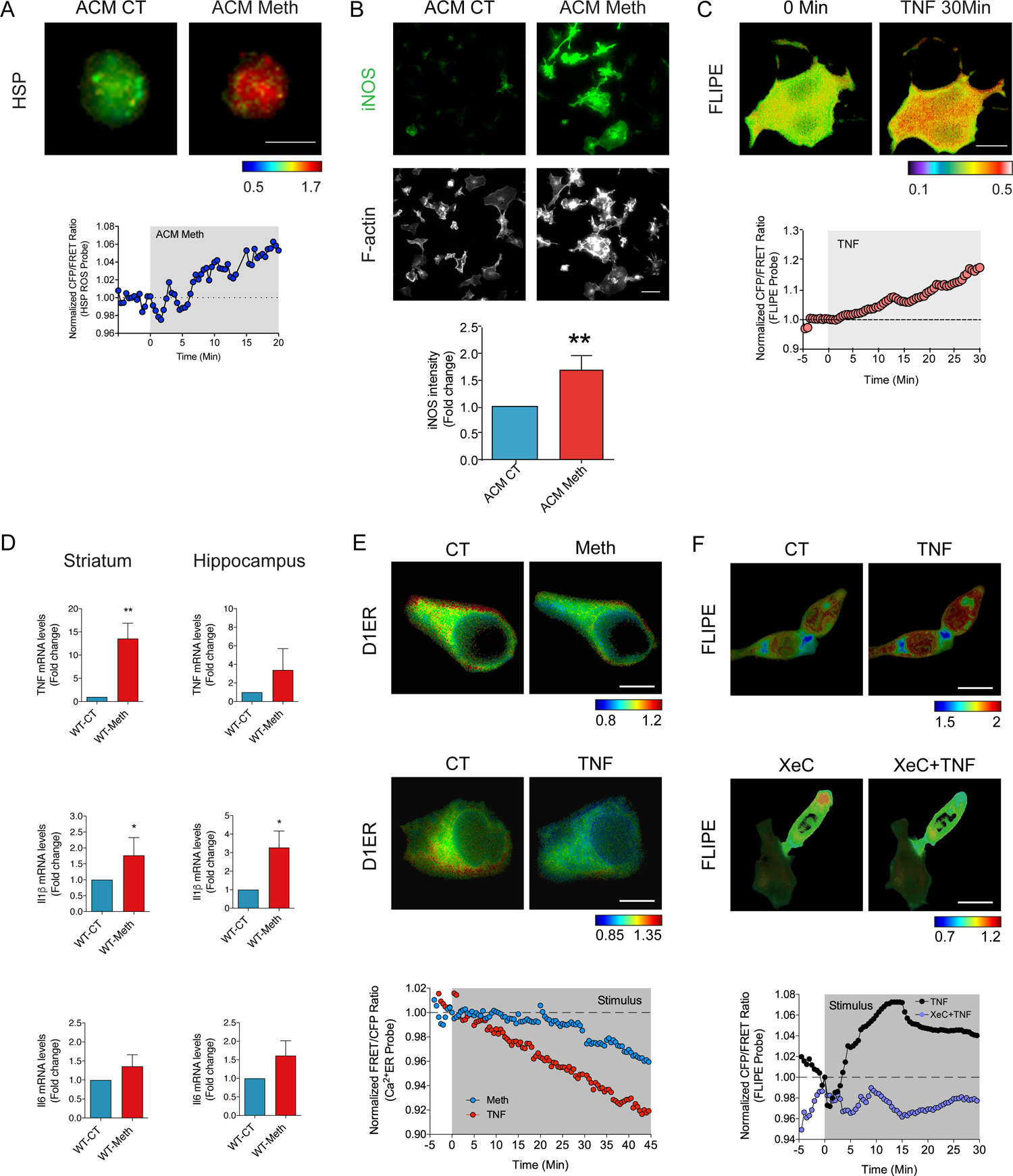
**A:** Primary cortical microglia expressing the ROS FRET biosensor (HSP) were incubated with ACM CT (left panel) and then exposed to ACM Meth (right panel). Time-lapses of CFP/FRET ratio changes for the HSP biosensor (normalized at 0 min) are shown according to the scale (n=4 cells pooled across two independent experiments). Scale bars, 10μm. **B:** Fluorescence imaging of primary cortical microglia immunolabeled for iNOS (green) and F-actin (grey; labeled with Alexa Fluor 647 Phalloidin obtained from Thermo Scientific (MA, USA)) and treated with ACM CT or ACM Meth for 24h (n=3 independent experiments). Graph (means and SEM) displays iNOS intensity normalized to the ACM CT. *p<0.05 (unpaired t test). Scale bar, 10μm. **C:** Primary cortical astrocytes expressing the glutamate release FRET biosensor (FLIPE) were exposed to TNF (50nM). Time-lapses of CFP/FRET ratio changes for the FLIPE biosensor (normalized at 0 min) are shown according to the scale (n=6 cells pooled across two independent experiments). Scale bars, 10μm. **D:** qRT-PCR for TNF, IL-1β and IL-6 from the striatum or hippocampus of mice administered with saline or binge Meth and sacrificed 24h after (n=4-5 mice *per* group). Graphs (means and SEM) display the fold change of indicated transcripts. *p<0.05 and **p<0.01 (unpaired t test). **E:** Primary cortical astrocytes expressing the endoplasmic reticulum calcium release FRET biosensor (D1ER) were exposed to Meth (100µM) (upper panels; blue circles) or TNF (50nM) (bottom panels; red circles). Time-lapses of CFP/FRET ratio changes for the D1ER biosensor (normalized at 0 min) are shown according to the scale (n=3-4 cells pooled across 2-3 independent experiments). Scale bars, 10μm. **F:** Primary cortical astrocytes expressing the glutamate release FRET biosensor (FLIPE) were exposed to TNF (50nM) (upper panels; black circles) or XestosponginC (500nM) + TNF (50nM) (bottom panels; lilac circles). Time-lapses of CFP/FRET ratio changes for the FLIPE biosensor (normalized at 0 min) are shown according to the scale (n=4 cells pooled across two independent experiments). Scale bars, 20μm.

**Supplementary Figure 4.**
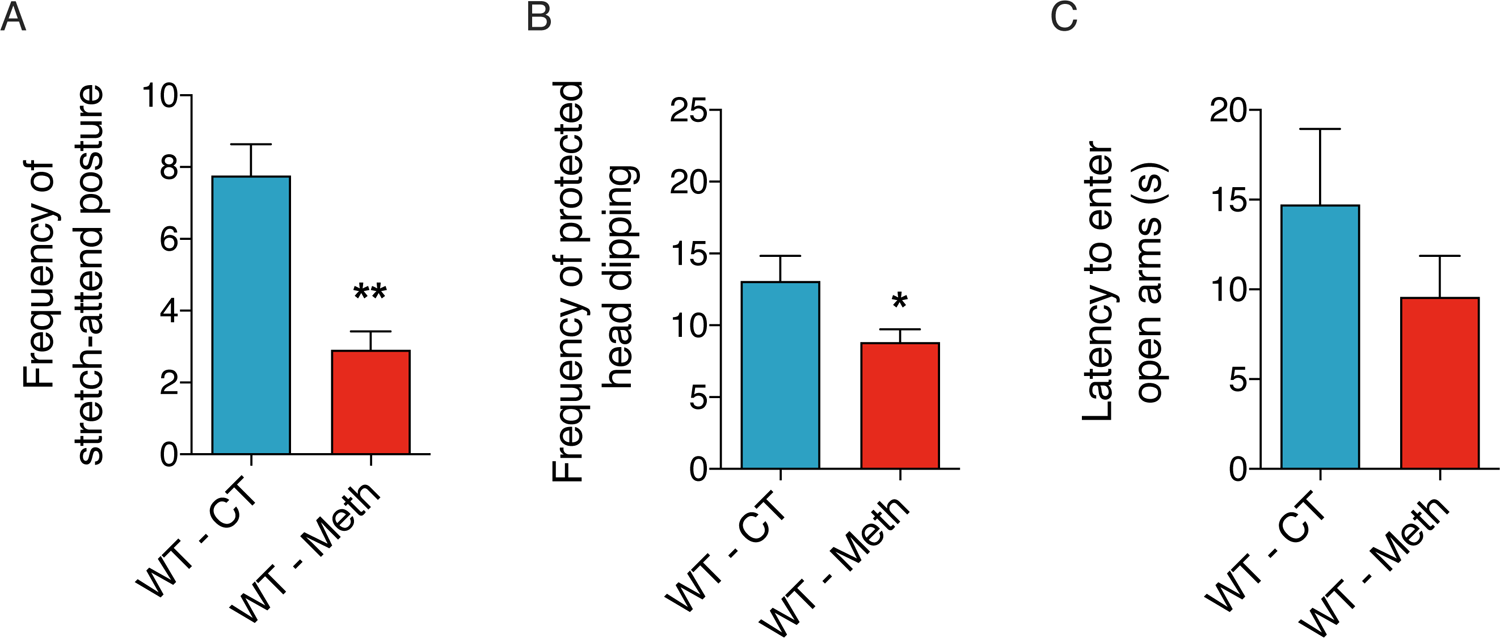
**A:** WT animals were evaluated in the EPM 24 hours after being administered with saline (CT) or binge Meth (n=11-13 animals *per* group). CT and Meth-treated mice displayed significant differences in the frequency of stretch-attend postures (SAP). The graph displays the mean and SEM. **p<0.01 (unpaired t test). **B:** WT animals were evaluated in the EPM 24 hours after being administered with saline (CT) or binge Meth (n=11-13 animals per group). CT and Meth-treated mice displayed significant differences in the frequency of protected head dipping. The graph displays the mean and SEM. *p<0.05 (unpaired t test). **C:** WT animals were evaluated in the EPM 24 hours after being administered with saline (CT) or binge Meth (n=11-13 animals per group). CT and Meth-treated mice displayed no differences regarding the latency to enter in open arms. The graph displays the mean and SEM.

**Supplementary Table 4.** Antibodies used for immunohistochemistry

| **Antibody** | **Dilution** | **Company** |
| --- | --- | --- |
| **GFAP** | 1:500 | Abcam (CAM, UK) |
| **Iba-1** | 1:500 | Wako (CA, USA) |
| **TNF** | 1:500 | Peprotech (LND, UK) |
| **Anti-mouse Alexa 488** | 1:1000 | Life Technologies (CA, USA) |
| **Anti-rabbit Alexa 568** | 1:1000 | Life Technologies (CA, USA) |

**Supplementary Table 5.** Antibodies used for immunocytochemistry

| **Antibody** | **Dilution** | **Company** |
| --- | --- | --- |
| **Arginase 1** | 1:100 | Santa Cruz Biotechnology (TX, USA) |
| **CD11b** | 1:200 | Abcam (CAM, UK) |
| **iNOS** | 1:200 | Santa Cruz Biotechnology (TX, USA) |
| **Anti-mouse Alexa 488** | 1:1000 | Life Technologies (CA, USA) |
| **Anti-rabbit Alexa 568** | 1:1000 | Life Technologies (CA, USA) |

**Supplementary Table 6.**
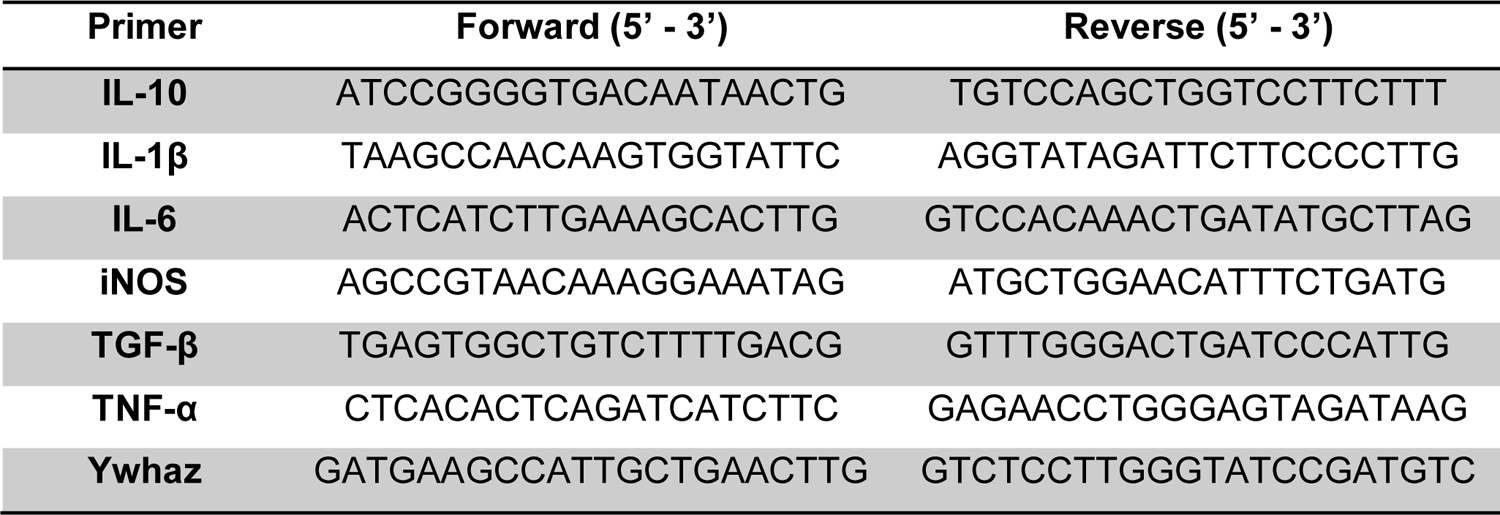
Primer sequences used in qRT-PCR

